# A cancer cell-intrinsic GOT2-PPARδ axis suppresses antitumor immunity

**DOI:** 10.1101/2020.12.25.424393

**Authors:** Hannah Sanford-Crane, Jaime Abrego, Chet Oon, Xu Xiao, Shanthi Nagarajan, Sohinee Bhattacharyya, Peter Tontonoz, Mara H. Sherman

## Abstract

Despite significant recent advances in precision medicine^1,2^, pancreatic ductal adenocarcinoma (PDAC) remains near-uniformly lethal. While the most frequent genomic alterations in PDAC are not presently druggable^3^ and conventional therapies are often ineffective in this disease^4^, immune-modulatory therapies hold promise to meaningfully improve outcomes for PDAC patients. Development of such therapies requires an improved understanding of the immune evasion mechanisms that characterize the PDAC microenvironment, including frequent exclusion of antineoplastic T cells and abundance of immune-suppressive myeloid cells^5–9^. Here we show that cancer cell-intrinsic glutamic-oxaloacetic transaminase 2 (GOT2) shapes the immune microenvironment to suppress antitumor immunity. Mechanistically, we find that GOT2 functions beyond its established role in the malate-aspartate shuttle^10–13^ and promotes the transcriptional activity of nuclear receptor peroxisome proliferator-activated receptor delta (PPARδ), facilitated by direct fatty acid binding. While GOT2 in PDAC cells is dispensable for cancer cell proliferation *in vivo*, GOT2 loss results in T cell-dependent suppression of tumor growth, and genetic or pharmacologic activation of PPARδ restores PDAC progression in the GOT2-null context. This cancer cell-intrinsic GOT2-PPARδ axis promotes spatial restriction of both CD4^+^ and CD8^+^ T cells from the tumor microenvironment, and fosters the immune-suppressive phenotype of tumor-infiltrating myeloid cells. Our results demonstrate a non-canonical function for an established mitochondrial enzyme in transcriptional regulation of immune evasion, which may be exploitable to promote a productive antitumor immune response.

## Main

Dual functions for GOT2 are described in the literature. The far better studied function is as a mitochondrial transaminase, implicated in maintenance of the malate-aspartate shuttle and redox homeostasis^10–13^. However, a limited body of evidence suggests a role for GOT2 in fatty acid binding and trafficking^14–19^, though this role remains poorly understood and has not been investigated in cancer. In these studies, GOT2 is often referred to as plasma membrane fatty acid binding protein (FABPpm) due to its membrane-proximal localization in hepatocytes and the ability of GOT2/FABPpm antiserum to disrupt fatty acid trafficking in metabolic cell types including hepatocytes and cardiomyocytes^14,20–22^. In light of recent work from our group and others documenting the importance of fatty acid trafficking for solid tumor progression^23–27^, we considered that GOT2 may promote PDAC growth, at least in part through its fatty acid trafficking function. GOT2 is overexpressed in human PDAC^28^, and while transmembrane fatty acid transporters were variably expressed, GOT2 was consistently expressed in human PDAC per two independent RNA-seq datasets (Extended Data Fig. 1a, 1b). We set out to determine whether GOT2 plays a role in PDAC progression *in vivo* and, if so, to understand the relevance of its established mitochondrial role versus its less characterized role in spatial regulation of fatty acids.

To assess the significance of GOT2 for PDAC progression, we generated several loss-of-function systems, using shRNA or CRISPR/Cas9 and using human and murine PDAC cells (Extended Data Fig. 2a). Cas9 and sgRNAs were introduced by transient transfection, and Cas9 was no longer expressed by the time cells were used for *in vivo* studies. Across all cell lines tested, only 2 showed proliferation defects (Fig. 1a, Extended Data Fig. 2b). These defects were modest and, in 1 of the 2 lines, a significant reduction in proliferation was only seen upon inducible GOT2 knockdown, suggesting that PDAC cells have sufficient metabolic plasticity to adapt to GOT2 loss and maintain proliferative capacity. However, when sgGot2 PDAC cells were transplanted into pancreata of immune-competent hosts, tumor growth was severely compromised (Fig. 1b). Consistent with *in vitro* results, proliferation among tumor cells was not impaired *in vivo* (Fig. 1c). An independent model also revealed a critical role for GOT2 in PDAC growth, whether GOT2 was knocked down with shRNA (Fig. 1d) or knocked out with CRISPR/Cas9 (Fig. 1e). Though shRNA-mediated knockdown had a less dramatic effect on tumor growth, we noted a partial recovery of GOT2 expression in these tumors by experimental endpoint (Extended Data Fig. 2c). These results suggested that GOT2 is dispensable for PDAC cell proliferation but required for tumor growth *in vivo*, and raised the possibility that cancer cell-intrinsic GOT2 promotes growth-permissive regulation of the tumor microenvironment.

**Figure 1:**
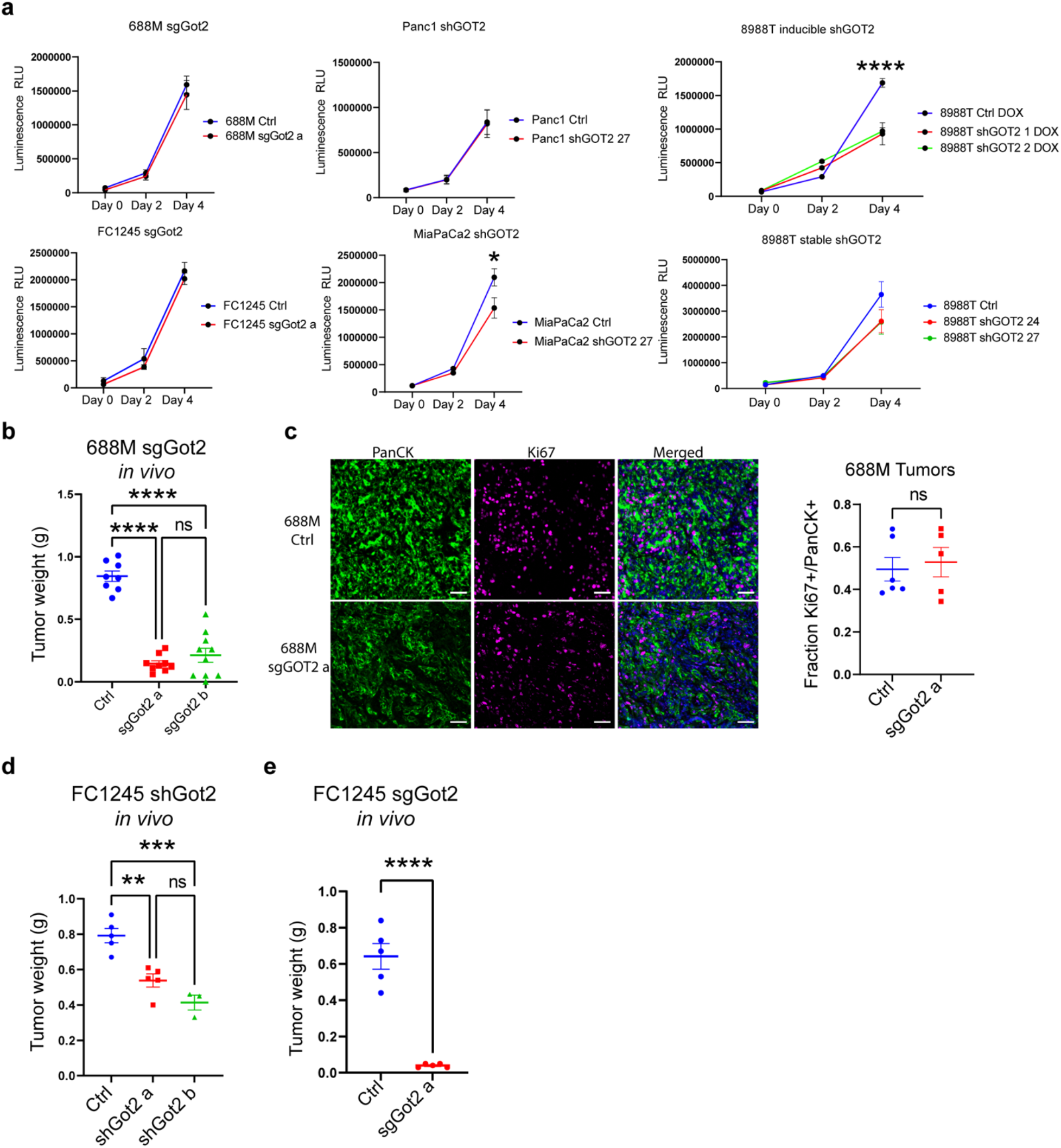
GOT2 promotes pancreatic tumor progression without impacting proliferation. **a,** Viable cell measurements in the indicated PDAC cell lines. Data are presented as mean ± s.e.m. from biological triplicates. *p < 0.05, ****p < 0.0001 by one-way ANOVA. **b,** PDAC tumor weight at experimental endpoint, 34 days after orthotopic transplantation of 688M cells into immune-competent hosts. Ctrl: n = 8, sgGot2 a: n = 9, sgGot2 b: n = 10. Data are presented as mean ± s.e.m. ****p < 0.0001 by one-way ANOVA. **c,** Immunohistochemical staining of tumors in **b** for Ki67 (proliferation) and pan-cytokeratin (panCK, tumor cells), with a DAPI counterstain (nuclei). Representative images are shown on the left (scale bar = 50 μm), with quantification on the right (Ctrl: n = 6, sgGot2: n = 5). Data are presented as mean ± s.e.m. ns = not significant by unpaired t-test. **d,** PDAC tumor weight at experimental endpoint, 22 days after orthotopic transplantation of FC1245 cells into immune-competent hosts. Ctrl: n = 5, shGot2 a: n = 5, shGot2 b: n = 3. Data are presented as mean ± s.e.m. **p < 0.01, ***p < 0.001 by one-way ANOVA. **e,** PDAC tumor weight at experimental endpoint, 18 days after orthotopic transplantation of FC1245 cells into immune-competent hosts. Ctrl: n = 5, sgGot2: n = 5. Data are presented as mean ± s.e.m. ****p < 0.0001 by unpaired t-test.

To gain insight into GOT2 function in an intact tumor microenvironment, we identified transcriptional programs with expression inversely correlated with *GOT2* transcript abundance in The Cancer Genome Atlas (TCGA) RNA-seq data (Fig. 2a). Pathway analysis of this group of genes revealed significant enrichment for genes associated with lymphocyte differentiation, activation, and adhesion, and led us to question whether cancer cell-intrinsic GOT2 regulates the abundance and/or activity of intratumoral T cells. We quantified T cells in 2 independent GOT2 loss-of-function models and found that T cell frequencies were significantly increased in sgGot2 or shGot2 tumors compared to controls, including CD4^+^ and CD8^+^ T cells (Fig. 2b-2d). As PDAC contains high numbers of immune-suppressive myeloid cells, including abundant macrophages, which contribute to T cell exclusion^29–31^, we assessed macrophage abundance and phenotype in these tumor tissues. We found that loss of GOT2 in cancer cells increased total macrophage abundance while decreasing the frequency of Arg1^+^ macrophages out of total macrophages (Fig. 2e), consistent with macrophage polarization to a less immune-suppressive phenotype permissive to T cell recruitment. The frequency of GRZB^+^ CD8^+^ cells was also increased in the sgGot2 setting (Fig. 2f, Extended Data Fig. 2d). To address whether the differences in T cell abundance were secondary to differences in tumor size, we performed a time course and harvested tumors soon after transplantation to quantify intratumoral T cells. At 11 days post-transplantation, a time point when tumors are small in control and sgGot2 tumors and not yet different in size, T cell frequencies are already significantly higher in the GOT2-null setting (Fig. 2g); T cell frequency was also significantly increased at days 19 and 27 post-transplantation, though tumors are significantly different in size by these early time points. We next asked whether these T cells were in fact functional in suppressing tumor progression. To address this, we treated control and sgGot2 tumors with neutralizing antibodies against CD4^+^ and CD8^+^ T cells. This intervention had no impact on growth of control tumors, consistent with previous studies documenting a lack of T cell-mediated antitumor immunity in mouse models of PDAC^32^. However, growth of sgGot2 tumors was restored upon T cell neutralization (Fig. 2h), suggesting that GOT2 promotes PDAC progression at least in part by suppressing T cell-dependent antitumor immunity.

**Figure 2:**
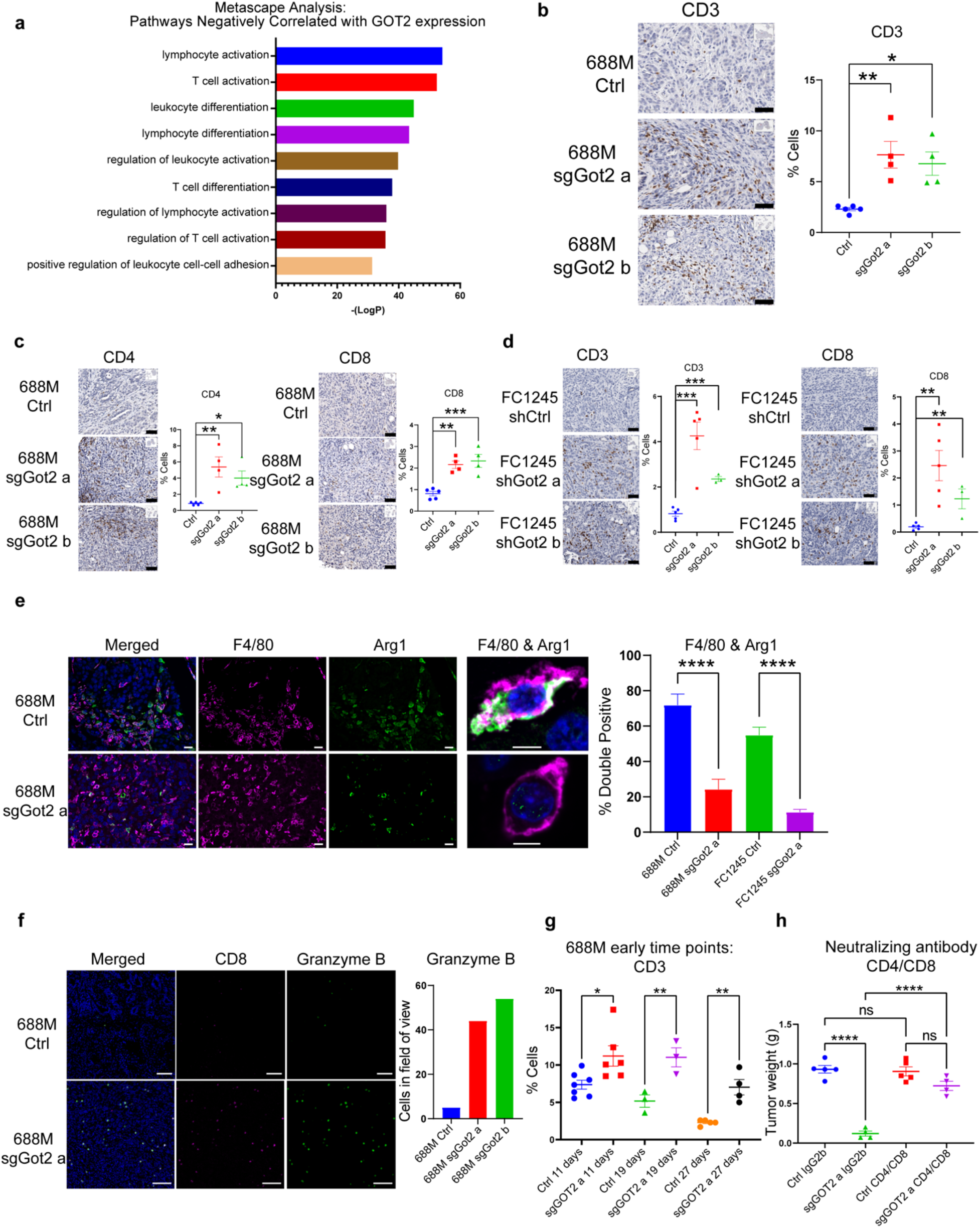
PDAC cell-intrinsic GOT2 suppresses T cell-dependent immunologic control of tumor growth. **a,** Metascape pathway analysis depicting transcriptional programs inversely correlated with GOT2 expression in human PDAC. **b,c,** Immunohistochemical staining of control and sgGot2 688M tumors for T cell marker CD3 (**b**) and subtype markers CD4 and CD8 (**c**). Representative images are shown on the left (scale bar = 50 μm), with quantification on the right (Ctrl: n = 5, sgGot2 a: n = 4, sgGot2 b: n = 4). **d,** Immunohistochemical staining of control and shGot2 FC1245 tumors for T cell markers CD3 and CD8. Representative images are shown on the left (scale bar = 50 μm), with quantification on the right (Ctrl: n = 5, shGot2 a: n = 5, shGot2 b: n = 3). **e,** Immunohistochemical co-staining of control and sgGot2 or shGot2 PDAC for macrophage marker F4/80 and immunosuppressive factor arginase-1. Representative images are from 688M tumors (scale bar on 20X images = 10 μm, scale bar on 63X images = 5 μm). Quantification of double-positive cells out of total F4/80^+^ cells in the 688M and FC1245 models is on the right; data are presented as mean ± s.e.m. ****p < 0.0001 by unpaired t-test. **f,** Immunohistochemical co-staining of control and sgGot2 688M tumors for T cell marker CD8 and granzyme B (scale bar = 50 μm), with granzyme B quantification on the right. **g,** Quantification of CD3 immunohistochemistry on 688M PDAC at the indicated time points post-transplantation (Ctrl d11: n = 7, sgGot2 d11: n = 6, Ctrl d19: n = 3, sgGot2 d19: n = 3, Ctrl d27: n = 5, sgGot2 d27: n = 4). *p < 0.05, **p < 0.01 by unpaired t-test. **h,** PDAC tumor weight at experimental endpoint, 27 days after orthotopic transplantation of 688M cells and treatment with isotype control or T cell neutralizing antibodies (details in Methods). Ctrl: n = 5 per cohort, sgGot2: n = 4 per cohort. For **b-h**, data are presented as mean ± s.e.m. For **b-e** & **h**, *p < 0.05, **p < 0.01, ***p < 0.001, ****p < 0.0001 by one-way ANOVA.

We next addressed the mechanism by which cancer cell-intrinsic GOT2 influences the immune microenvironment, taking potential enzymatic and fatty acid-binding functions into account. To begin to address this, we examined GOT2 localization in PDAC cells and found that a pool of this canonically mitochondrial protein localizes to the nucleus in murine pre-malignant lesions and PDAC as well as human PDAC *in vivo* (Fig. 3a, 3b, Extended Data Fig. 3a). We note that, while all human PDAC specimens examined showed evidence of nuclear GOT2 in pan-cytokeratin^+^ tumor cells, tumor cells with GOT2 restricted to mitochondrial and membrane-proximal regions and without nuclear GOT2 were also observed across these samples. This nuclear GOT2 pool was also evident *in vitro*, whether we analyzed endogenous or exogenous, His-tagged GOT2 (Extended Data Fig. 3b-e). We reasoned that the intact proliferation of GOT2-null tumors suggested the presence of metabolic adaptation mechanisms to retain redox balance, and motivated us to consider non-canonical functions of GOT2 related to its putative fatty acid binding capacity. The previously unappreciated nuclear pool of GOT2 led us to hypothesize that GOT2 regulates nuclear trafficking of fatty acids, either into or within the nucleus. Nuclear fatty acid trafficking has been shown to be regulated by fatty acid binding proteins^33,34^, and nuclear fatty acids have established functional significance as ligands for the peroxisome proliferator-activated receptor (PPAR) members of the nuclear receptor superfamily of transcription factors^35^. This 3-member family is activated by fatty acid ligands, and while PPARα and PPARγ display tissue-restricted expression, PPARδ is ubiquitously expressed, and was expressed in all PDAC lines examined, whether or not GOT2 was inhibited (Extended Data Fig. 4a). Importantly, PPARδ promotes tumorigenesis via tissue-specific metabolic and immune-modulatory mechanisms^36–39^, prompting us to test a functional relationship between GOT2 and PPARδ that may underlie the phenotypes of GOT2-null PDAC.

**Figure 3:**
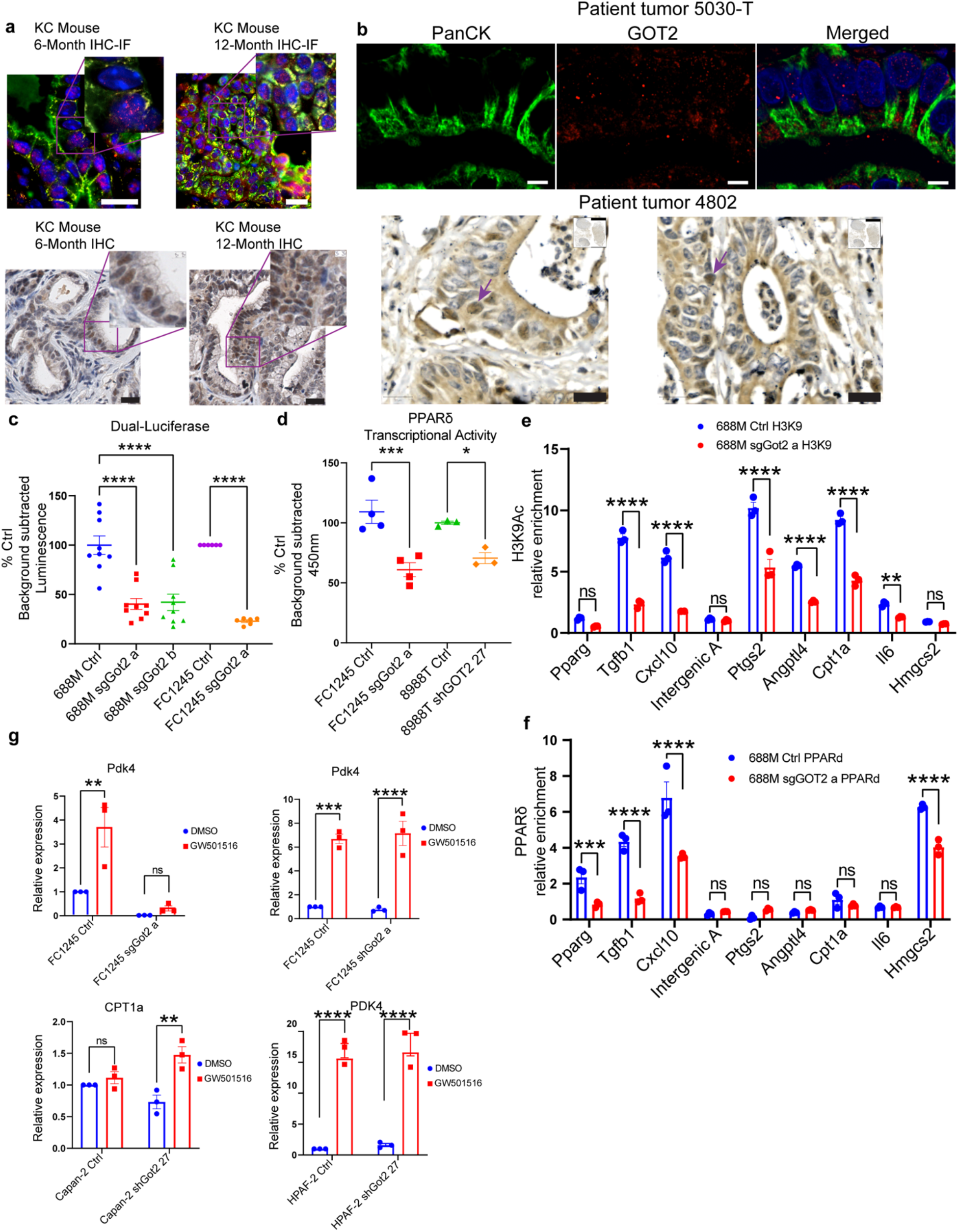
GOT2 positively regulates PPARδ activity. **a,** Immunohistochemical staining for GOT2 or GOT2 and panCK in pancreas tissues from *Kras^LSL-G12D/+^;Pdx1-Cre* (KC) mice at 6 or 12 months of age (representative of n = 4 per time point). Scale bar = 20 μm. **b,** Immunohistochemical staining for GOT2 or GOT2 and panCK in human PDAC (representative of n = 5). Fluorescent images: scale bar = 5 μm, brightfield image: scale bar = 20 μm. Arrows indicate examples of tumor cells with nuclear GOT2. **c,** Luciferase assay for PPRE activity in the indicated cell lines, normalized to renilla, presented as mean ± s.e.m. ****p < 0.0001 by one-way ANOVA (688M) or unpaired t-test (FC1245). **d,** PPARδ transcriptional activity assay, reading out binding to immobilized DNA containing PPREs, in the indicated cell lines. Data are presented as mean ± s.e.m. from 4 (FC1245) or 3 (8988T) independent experiments. *p < 0.05, ***p < 0.001 by unpaired t-test. **e,f,** Chromatin immunoprecipitation (ChIP) for (**e**) H3K9Ac and (**f**) PPARδ in control or sgGot2 688M PDAC cells, followed by qPCR for proximal promoter regions of the indicated genes. Data were normalized to an intergenic region (intergenic B; intergenic A was an additional control region) and are presented as mean ± s.e.m. from biological triplicates. **p < 0.01, ***p < 0.001 ****p < 0.0001 by unpaired t-test. **g,** qPCR for the indicated PPARδ-regulated genes in control or GOT2-knockdown PDAC cells, treated with vehicle (DMSO) or PPARδ synthetic agonist GW501516 (100 nM). Data are presented as mean ± s.e.m. from biological triplicates. **p < 0.01, ***p < 0.001, ****p < 0.0001 by unpaired t-test.

Transcriptional activity from a PPAR response element (PPRE) was reduced in GOT2-null PDAC cells (Fig. 3c), suggesting that GOT2 positively regulates PPARδ activity. Unlike steroid-activated nuclear receptors, which are sequestered in the cytoplasm in the absence of ligand and translocate to the nucleus upon ligand engagement, PPARδ is constitutively nuclear and bound to DNA, but undergoes a conformational change upon binding of nuclear fatty acids to enable interaction with coactivator complexes, altered DNA binding, and induction of target gene expression^40^. Further supporting positive regulation of PPARδ transcriptional activity by GOT2, nuclear extracts from control and sgGot2 cells were applied to wells containing immobilized, PPRE-containing DNA, followed by incubation with a PPARδ antibody and a peroxidase-conjugated secondary antibody. Results of this assay suggested reduced PPARδ transcriptional activity in GOT2-null PDAC cells (Fig. 3d). Chromatin immunoprecipitation (ChIP) PPARδ and acetylated histone H3K9, a marker of active promoters, followed by qPCR also supported a reduction of PPARδ transcriptional activity in the absence of GOT2 (Fig. 3e, 3f). Some of these genes previously linked to PPARδ activity appear potentially to be indirect targets. Expression of PPARδ target genes was also reduced in GOT2-null PDAC cells, and synthetic PPARδ agonist GW501516 restored target gene expression, suggesting that these genes are indeed PPARδ-regulated (Fig. 3g). Among the genes with clear relevance to our *in vivo* phenotype was *PTGS2*, which encodes COX-2. Recently reported gain- and loss-of-function experiments suggest that COX-2 promotes T cell exclusion from the PDAC microenvironment, consistent with our results, and that *PTGS2* expression correlated with poor patient survival^41^. We further investigated regulation of COX-2 downstream of GOT2 and found that COX-2 protein levels were reduced in GOT2-null PDAC cells *in vitro* and *in vivo* (Extended Data Fig. 4a, 4b). These results together suggest that GOT2 promotes transcriptional activity of PPARδ in PDAC cells.

As we were prompted to investigate a GOT2-PPARδ functional interaction based on the putative fatty acid binding function of GOT2, we investigated this role further. For this, we analyzed the crystal structure of human GOT2^42^ and identified 5 putative fatty acid binding sites based on hydrophobicity (Fig. 4a, 4b). We then performed *in silico* docking studies for known fatty acid ligands for PPARδ, and identified a potential interaction between arachidonic acid and GOT2 hydrophobic site 2 (Fig. 4c). This modeled interaction yielded a docking score of −7.6 kcal/mol, which is very similar to the docking score calculated for arachidonic acid in the ligand binding domain of PPARγ (−7.0 kcal/mol)^43^, an interaction that is known to be direct and functionally significant. To probe this interaction further, we performed competitive fatty acid binding assays using purified GOT2 protein and radiolabeled arachidonic acid. In addition to cold arachidonic acid, we used cold oleic acid as this was previously suggested to bind to GOT2^19^ (though our analysis revealed a distinct fatty acid binding domain than this previous study) as well as prostaglandin D2 (PGD_2_), a downstream metabolite of arachidonic acid which we predicted to serve as a negative control and not to bind directly to GOT2 based on computational modeling. The competitive binding assay showed that arachidonic acid indeed bound to GOT2 directly, and while cold arachidonic acid readily displaced radiolabeled ligand, our negative control lipid (PGD_2_) was unable to compete away the arachidonic acid signal even when PGD_2_ concentration exceeded that of arachidonic acid by three orders of magnitude (Fig. 4d), supporting a specific interaction. Oleic acid had a modest effect on binding, suggesting that oleic acid may bind to the arachidonic acid-bound site but at a lower affinity, or may bind to a separate site on the protein. To assess a relationship between GOT2 and arachidonic acid trafficking in cells, we performed mass spectrometry to measure arachidonic acid in whole cells and nuclei; levels were unchanged at the whole-cell level between control and sgGot2 cells (Extended Data Fig. 4c), but nuclear levels were below a reliable level of detection. We developed an assay to measure nuclear arachidonic acid accumulation by spiking fluorescent arachidonic acid into our culture medium and measuring fluorescent signal in isolated nuclei, which revealed a significant reduction in nuclear arachidonic acid accumulation in two GOT2 loss-of-function cell lines (Fig. 4e). Though significant, these differences were modest, suggesting that GOT2 may regulate arachidonic acid within the nucleus as opposed to predominantly regulating its nuclear import. To address the functional significance of GOT2 fatty acid binding, we looked closely at the putative fatty acid binding pocket we identified, and selected 3 key amino acid residues we predicted to be critical for arachidonic acid binding at that site (Fig. 4f). We mutated these 3 residues on His-tagged GOT2 and used this triple-mutant GOT2 (tmGOT2) or wild-type GOT2 (wtGOT2) to reconstitute sgGot2 PDAC cells. While wtGOT2 localized to mitochondria and nuclei, tmGOT2 showed a preferential mitochondrial localization compared to the wild-type protein (Fig. 4g-4i), raising the possibility that fatty acid binding at this site promotes GOT2 nuclear trafficking, perhaps via interaction with a chaperone. After confirming that tmGOT2 retains enzymatic activity (Extended Data Fig. 5a, 5b), we assessed PPARδ activity and found that target gene expression and transcriptional activity were reduced in cells expressing tmGOT2 compared to wtGOT2 (Fig. 4j, 4k). We next transplanted immune-competent mice with control, sgGot2, sgGot2 + wtGot2, or sgGot2 + tmGot2 PDAC cells. While wtGot2 completely rescued tumor growth as expected, tmGot2 only partially rescued tumor growth (Fig. 4l), suggesting a significant role for this fatty acid binding region in GOT2-mediated PDAC progression.

**Figure 4:**
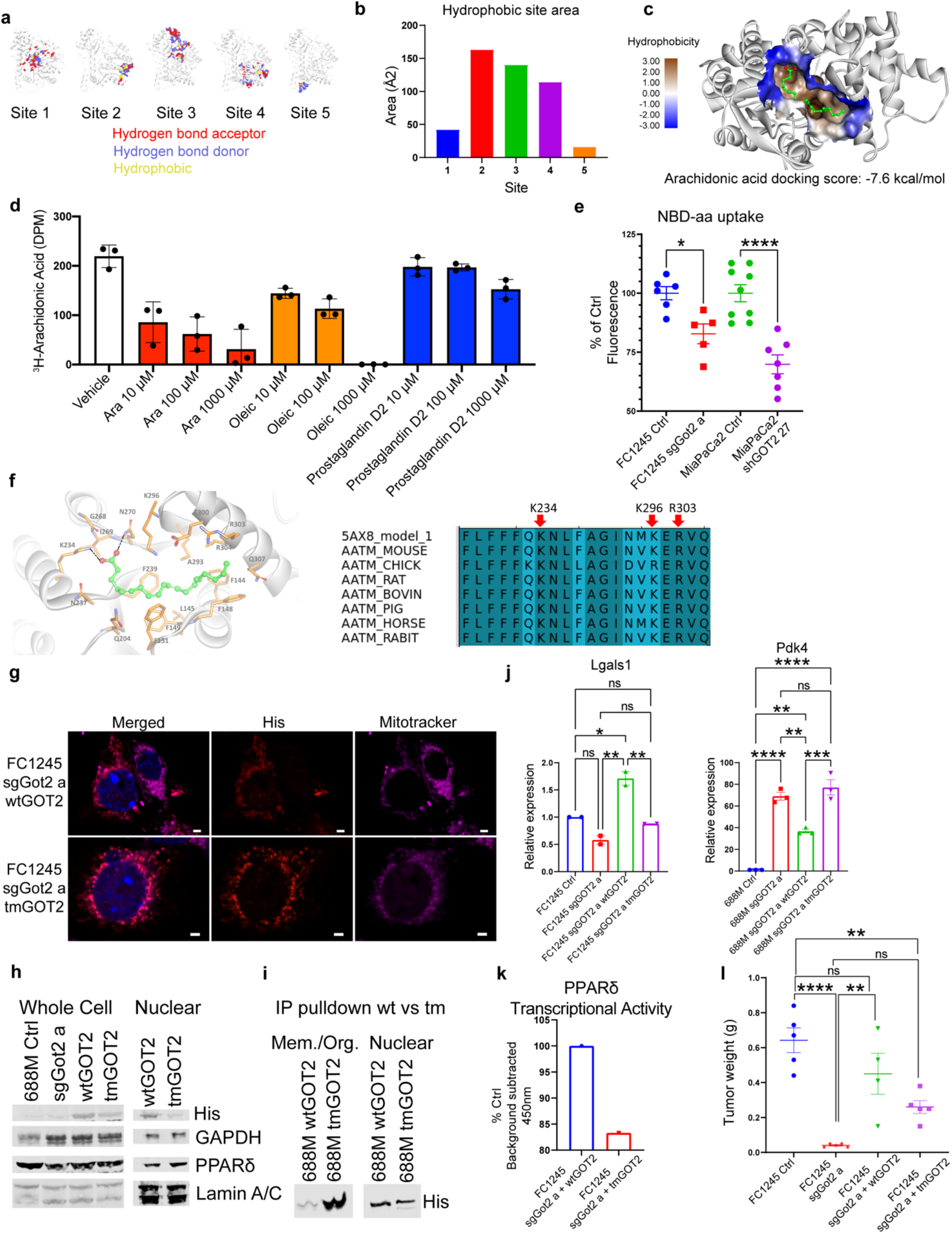
GOT2 binds to PPARδ ligand directly. **a,** Hydrophobic site maps on the GOT2 protein, indicating putative fatty acid binding domains. Red: hydrogen bond acceptor, blue: hydrogen bond donor, yellow: hydrophobic. **b,** Plot of the hydrophobic area of the putative fatty acid binding sites depicted in **a**. **c,** Docking model of arachidonic acid in site 2 on the GOT2 protein, with bioenergetic docking score (−7.6 kcal/mol) indicated below. **d,** Competitive fatty acid binding assay, measuring radioactivity upon incubating purified human GOT2 with ^3^H-arachidonic acid (1 μM) and the indicated concentrations of cold lipid species. **e,** Nuclear accumulation of NBD-arachidonic acid after the indicated cell lines were incubated with 2.5 μM NBD-aa for 2 hr (MiaPaCa2) or 2 μM NBD-aa for 15 min (FC1245). Data are presented as mean ± s.e.m. *p < 0.05, ****p < 0.0001 by unpaired t-test. **f,** (Left) Model of arachidonic acid bound to GOT2, indicating amino acid residues that potentially facilitate binding. Based on this model, K234, K296, and R303 were selected for mutation to alanine. (Right) Conservation of GOT2 amino acid sequence, including the 3 residues predicted to support arachidonic acid binding, among higher vertebrates. **g,** Immunofluorescence staining of wtGOT2 and tmGOT2 (both His-tagged) in PDAC cells, with a DAPI nuclear counterstain. Scale bar = 2 μm. **h,** Western blot on whole cell lysates and nuclear extracts on the indicated 688M stable cell lines. **i,** His immunoprecipitation and Western blot from membrane/organelle fractions versus nuclear fractions in the indicated 688M stable cell lines. **j,** qPCR for the indicated PPARδ-regulated genes in FC1245 stable cell lines, normalized to *36b4*. Data are presented as mean ± s.e.m. from biological triplicates. *p < 0.05, **p < 0.01, ***p < 0.001, ****p < 0.0001 by one-way ANOVA. **k,** PPARδ transcriptional activity assay in the indicated FC1245 stable cell lines. **l,** PDAC tumor weight at experimental endpoint, 18 days after orthotopic transplantation of the indicated FC1245 cells. Ctrl: n = 5, sgGot2: n = 5, sgGot2 + wtGOT2: n = 4, sgGot2 + tmGOT2: n = 5. Ctrl and sgGot2 arms here are also depicted in Figure 1e. Data are presented as mean ± s.e.m. **p < 0.01, ****p < 0.0001 by one-way ANOVA.

Based on these results, we hypothesized that PPARδ activation would restore PDAC growth in the GOT2-null setting. We treated PDAC cells with GW501516 as we found this to override the limitation on PPARδ activity in sgGot2 cells *in vitro*, and observed no increase (in fact, a decrease) in proliferation (Fig. 5a). However, GW501516 treatment *in vivo* rescued growth of GOT2-null PDAC without impacting control tumor growth (Fig. 5b), and restored immune suppression with respect to intratumoral T cell abundance (Fig. 5c) and induction of COX-2 expression (Fig. 5d, Extended Data Fig. 4b). As GW501516 acts systemically, we next specifically activated PPARδin PDAC cells by introducing a fusion of PPARδ with the VP16 transactivation domain from herpes simplex virus^44^, to enable ligand-independent activation, into control and sgGot2 PDAC cells (Extended Data Fig. 5c) at sufficiently low copy number to avoid detectable PPARδ overexpression. While VP16-PPARδ did not increase proliferation *in vitro* (Fig. 5e) nor increase PDAC growth in the control group, genetic PPARδ activation significantly albeit partially rescued tumor growth in sgGot2 tumors in 2 independent models (Fig. 5f, 5g). Consistent with these findings, VP16-PPARδ increased expression of target genes such as *Ptgs2* in sgGot2 cells (Fig. 5h). Together, these results suggest that GOT2 promotes PDAC progression and immune suppression by activating PPARδ.

**Figure 5:**
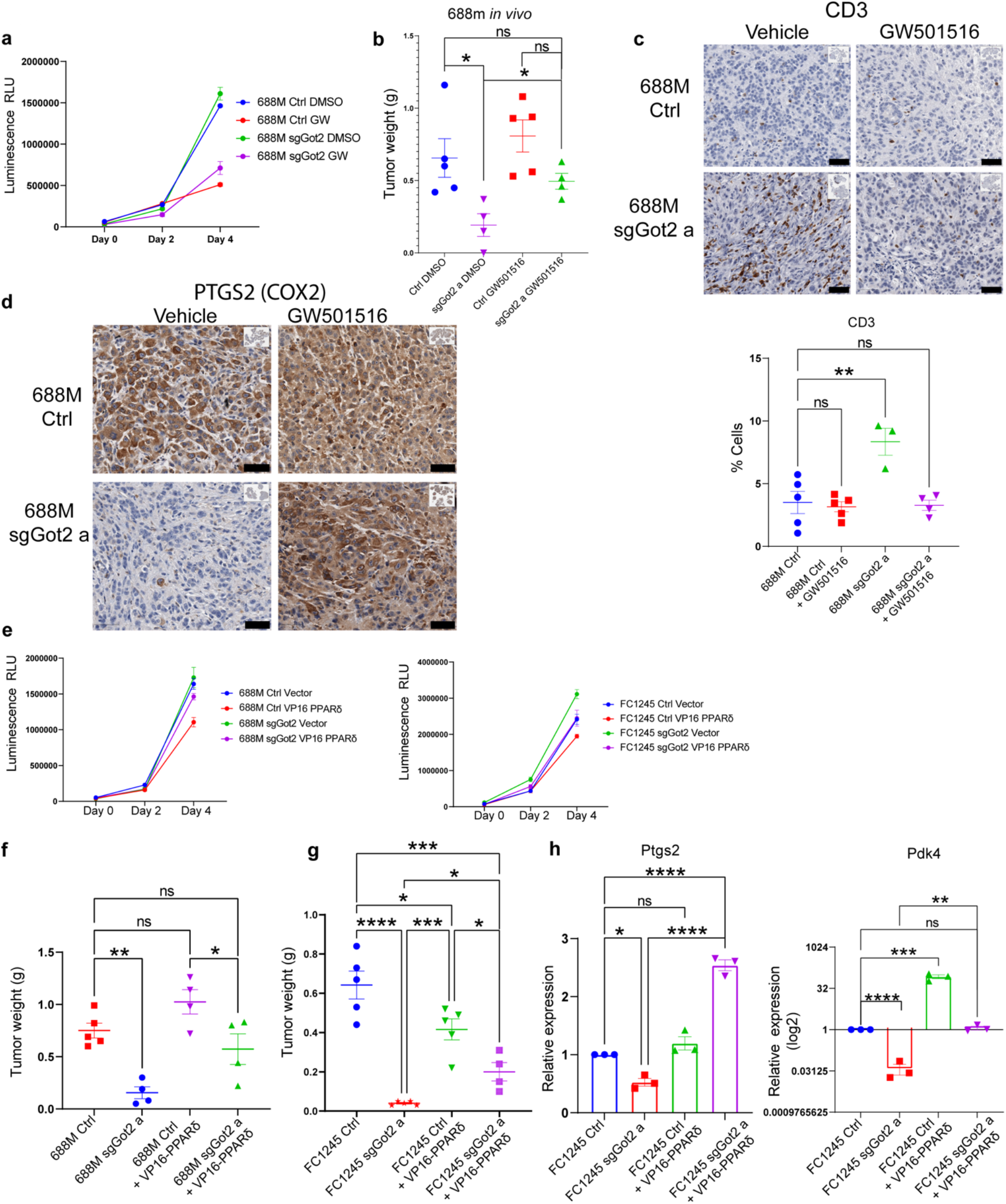
PPARδ activation restores tumor growth and T cell exclusion in the absence of GOT2. **a,** Viable cell measurements in control or sgGot2 PDAC cells treated with vehicle or 100 nM GW501516. **b,** PDAC tumor weight at experimental endpoint, 30 days after orthotopic transplantation of the control or sgGot2 cells, with daily i.p. injection of vehicle or 4 mg/kg GW501516. Ctrl: n = 5 per cohort, sgGot2: n = 4 per cohort. Data are presented as mean ± s.e.m. *p < 0.05 by one-way ANOVA. **c,** Immunohistochemical staining of control and sgGot2 688M tumors treated with vehicle or GW501516 as in **b** for T cell marker CD3. Representative images are shown above (scale bar = 50 μm), with quantification below (Ctrl: n = 5, Ctrl + GW501516: n = 5, sgGot2: n = 3, sgGot2 + GW501516: n = 4). Data are presented as mean ± s.e.m. **p < 0.01 by one-way ANOVA. **d,** Immunohistochemical staining for PTGS2/COX2 in control or sgGot2 PDAC treated with vehicle or GW501516 (representative of n = 3-5 per cohort). Scale bar = 50 μm. **e,** Viable cell measurements in control or sgGot2 PDAC cells stably transduced with empty vector or VP16-PPARδ. Data are presented as mean ± s.e.m. **f,g,** PDAC tumor weight at experimental endpoint in the indicated (**f**) 688M and (**g**) FC1245 lines. 688M: Ctrl: n = 5, sgGot2: n = 4, Ctrl VP16-PPARδ: n = 4, sgGot2 VP16-PPARδ: n = 4, endpoint = day 27. FC1245: Ctrl: n = 5, sgGot2: n = 5, Ctrl VP16-PPARδ: n = 5, sgGot2 VP16-PPARδ: n = 4, endpoint = day 18. Ctrl and sgGot2 FC1245 arms here are also depicted in Figure 1e. *p < 0.05, **p < 0.01, ***p < 0.001, ****p < 0.0001 by one-way ANOVA. **h,** qPCR for PPARδ-regulated genes in the indicated FC1245 stable cell lines, normalized to *36b4.* Data are presented as mean ± s.e.m. from biological triplicates. *p < 0.05, **p < 0.01, ***p < 0.001, ****p < 0.0001 by one-way ANOVA.

Our study suggests that GOT2 plays a critical role in promoting a tumor-permissive immune microenvironment in the pancreas. This function is attributable at least in part to direct fatty acid binding, and activation of nuclear receptor PPARδ. Further studies are needed to understand the mechanisms regulating GOT2 subcellular localization, as well as the precise molecular mechanism by which GOT2 promotes PPARδ transcriptional activity. While diverse mechanisms contribute to immune evasion in PDAC^45^, targeting GOT2 may be part of a potential treatment approach to foster an immune response against this deadly cancer.

## Methods

### Animals

All experiments were reviewed and overseen by the institutional animal use and care committee at Oregon Health and Science University in accordance with NIH guidelines for the humane treatment of animals. C57BL/6J (000664, for models with FC1245^1^) or B6129SF1/J (101043, for models with 688M^2^) mice from Jackson Laboratory were used for orthotopic transplant experiments at 8-10 weeks of age. Tissues from 6- or 12-month-old *Kras^LSL-G12D/+^;Pdx1-Cre* (KC) mice were kindly provided by Dr. Ellen Langer (OHSU).

### Human tissue samples

Human patient PDAC tissue samples donated to the Oregon Pancreas Tissue Registry program (OPTR) in accordance with full ethical approval were kindly shared by Dr. Jason Link and Dr. Rosalie Sears (OHSU).

### Plasmids

The pCMX-VP16-PPARD plasmid was kindly provided by Dr. Vihang Narkar (University of Texas Health Science Center at Houston)^3^. The VP16-PPARD element was cloned into the lentiviral vector. To construct pLenti VP16 PPARD, the VP16-PPARD element was amplified by PCR using sense primer 5’-GGGGACAAGTTTGTACAAAAAAGCAGGCTTAATGGCCCCCCCGAC-3’ and antisense primer 5’-GGGGACCACTTTGTACAAGAAAGCTGGGTTTTAGTACATGTCCTTGTAGATTTCCTGGAGCAGG-3’. PCR product was inserted into pDONR 221 entry clone using Gateway BP Clonase II enzyme (Thermo Fisher 12535029). Entry clone VP16 PPARD element was swapped into expression region of pLenti CMV Puro DEST (Addgene #17452) using LR Clonase II enzyme (Thermo Fisher 11-791-020) to generate pLenti VP16 PPARD construct. The pCMV3 plasmid containing C-terminal His-tagged human GOT2 cDNA was purchased from Sino Biological (HG14463-CH) and cloned into the lentiviral vector pLenti CMV Puro DEST (Addgene #17452) using the same approach as pLenti-VP16 PPARD. pLenti wtGOT2 PCR product was generated using sense primer 5’-GGGGACAAGTTTGTACAAAAAAGCAGGCTTAATGGCCCTGCTGCACT-3’ and antisense primer 5’-GGGGACCACTTTGTACAAGAAAGCTGGGTTTTTAGTGATGGTGGTGATGATGGTGG-3’. Triple mutant GOT2 was constructed using Q5 Site-Directed Mutagenesis Kit (New England E0552S) in two subsequent steps. Two sets of primers were used to generate three site mutations; primer set 1 for K234A mutation (F:5’-AACAGTGGTGGCGAAAAGGAATCTC-3’; R:5’-GCTATTTCCTTCCACTGTTC-3’) and primer set 2 for K296A and R303A mutations (F:5’-GTCTGCGCAGATGCGGATGAAGCCAAAGCGGTAGAGTC-3’; R:5’-CATAGTGAAGGCTCCTACACGC-3’). pLenti tmGOT2 was then generated using the same approach and primers as pLenti wtGOT2.

### Cell lines

Human pancreatic cancer cell lines MIAPaCa-1, PA-TU-8988T, Panc1, HPAF-II, and Capan-2 were obtained from ATCC and grown in Dulbecco’s modified Eagle’s medium (DMEM) containing 10% fetal bovine serum. Non-transformed, TERT-immortalized human pancreatic ductal epithelial cells were kindly provided by Dr. Rosalie Sears (OHSU)^4^. PA-TU-8988T cells harboring doxycycline-inducible shGOT2 were kindly provided by Dr. Costas Lyssiotis (University of Michigan). FC1245 PDAC cells were generated from a primary tumor in *Kras^LSL-G12D/+^;Trp53^LSL-R172H/+^;Pdx1-Cre* mice and were kindly provided by Dr. David Tuveson (Cold Spring Harbor Laboratory)^1^. 688M PDAC cells were generated from a liver metastasis in *Kras^LSL-G12D/+^;Trp53^LSL-R172H/+^;Pdx1-Cre;Rosa26^LSL-tdTomato/+^* mice and were kindly provided by Dr. Monte Winslow (Stanford University School of Medicine)^2^. Cell lines were routinely tested for *Mycoplasma* at least monthly (MycoAlert Detection Kit, Lonza).

The pSpCas9(BB)-2A-Puro(PX459) v2.0 plasmid (Addgene #62988) was used to clone guide sequences targeting Got2 per supplier’s protocol; sgRNA A: GACGCGGGTCCACGCCGGT, sgRNA B: ACGCGGGTCCACGCCGGTG. The 688M or FC1245 cell line was transfected with control plasmid or plasmid containing either of the sgGot2 sequences and subject to selection with 2 μg/ml puromycin for 4 days. Single-cell clones were expanded and screened for GOT2 protein expression by Western blot.

GOT2 shRNA vectors were purchased in bacterial glycerol stocks from Sigma-Aldrich Mission shRNA (mouse shRNA A: TRCN0000325948, shRNA B: TRCN00000325946; human sh24: TRCN0000034824, sh27: TRCN0000034827). Briefly, bacterial cultures were amplified in ampicillin growth medium from glycerol stocks for use in purification of plasmid DNA. Subsequently, purified plasmid was transfected to packaging cells HEK293T for production of lentiviral particles. FC1245 cells were then infected and puromycin selected to generate stable GOT2 knockdowns, with validation by Western blot. Lentivirus preparation for stable cell line generation was done with pMD2.G envelope plasmid (Addgene #12259) and psPAX2 packaging plasmid (Addgene 12260) in 293T-LentiX cells. Briefly, 5 μg pMD2.G, 5 μg psPAX2 and 10 μg of plasmid DNA (shGOT2 KD, VP16-PPARdelta, wtGOT2, tmGOT2, or scramble Ctrl) were combined with 600 μl Opti-MEM and 20 μl lipofectamine 2000 for 20 mins at room temp. 10cm dishes of 293T-LentiX were kept in 0% FBS DMEM and the mixture was added in a dropwise manner. 12hrs later media was changed to 10% FBS DMEM. At 24hrs after transduction and 48hrs after transduction, media was collected and filtered through a 0.25 μm filter, aliquoted, and frozen at −80°C. Lentiviral transduction of human and mouse cell lines: cells were plated to 6-well plates. 10 μg/mL polybrene (EMD Millipore TR-1003-G) was added to 1 mL 10% FBS DMEM and 300 μl of filtered lentivirus media. 24hrs later media was changed to fresh 10% FBS DMEM. 48hrs after initial transduction, cells were treated with 2 μg/mL puromycin (Thermo Fisher A1113803), or 4 μg/mL puromycin depending on cell line. A control well of non-transduced cells was used as an indicator for proper selection. Protein knockdown was validated by Western blot.

### Western Blotting

PDAC cells were treated as described in the text, and whole cell lysates were prepared in RIPA buffer containing protease inhibitor cocktail (Sigma-Aldrich 11836170001). Alternatively, sub-cellular fractions were prepared using Cell Fractionation Kit #9038 purchased from Cell Signaling Technology following the manufacturer’s instructions. Briefly, cells were collected with scraping, washed in PBS and pelleted (350 x g 5mins). Cells were resuspended in 500 μl PBS and 100 μl reserved for whole cell lysis in RIPA buffer + cOmplete mini EDTA-free protease inhibitor cocktail. Remaining cell pellet was centrifuged (500 x g 5 mins), PBS was decanted and 500 μl CIB + 5 μl Protease Inhibitor and 2.5ul PMSF was added. Solutions were vortexed and stored on ice for 5 mins. Lysates were centrifuged (500 x g 5mins); supernatant was collected as the cytosolic fraction. Remaining insoluble pellet was washed with CIB and supernatant decanted. 500 μl MIB + 5ul Protease Inhibitor and 2.5ul PMSF was then added to the cell pellet. After vortexing 15 seconds, solutions were incubated on ice for 5 mins, and centrifuged (8000 x g 5 mins). Supernatant was collected as the membrane & organelle fraction. Pellet was then washed in MIB and supernatant decanted. 250 μl CyNIB + 2.5 μl Protease Inhibitor + 1.25 μl PMSF was then added to the pellet containing nuclei. Solution was sonicated for 5 sec at 20% power 3x to prepare nuclear lysate. For western blot, 60 μl 4X LDS loading buffer with 10X reducing agent was added for every 100 μl of supernatant per fraction. Samples were boiled for 5 mins at 95C and centrifuged for 3 mins at 15,000 x g. 15 μl of each fraction along with 15 μl of whole cell lysate was loaded for Western blotting. Alternatively, to generate total nuclear and cytosolic fractions, NE-PER Nuclear and Cytoplasmic Extraction Reagent Kit (Thermo Fisher) was used according to the manufacturer’s protocol. Where indicated, His-tagged GOT2 protein was immunoprecipitated using the His-tag isolation and pull-down Dynabeads system (Thermo Fisher) using the manufacturer’s protocol. Protein concentration was quantitated using the BCA protein assay kit (Pierce). Equal amounts of protein were loaded in each lane and separated on a 4-12% Bis-Tris NuPAGE gel (Invitrogen), then transferred onto a PVDF membrane. Membranes were probed with primary antibodies and infrared secondary antibodies: GOT2 (Thermo Fisher PA5-77990), β-Actin (Santa Cruz sc-47778), PPARδ (Abcam ab178866), His (R&D Systems MAB050-100), COX2 (Abcam ab15191), COX IV (Cell Signaling Technology 11967S), AIF (Abcam ab1998), PMCA1 (Novus Biologicals 5F10), Lamin A/C (Cell Signaling Technology 4777S), Tom20 D8T4N (Cell Signaling Technology 42606S), HSC 70 (Santa Cruz sc-7298), anti-rabbit Alexa Fluor Plus 680 (Thermo Fisher A32734) and anti-mouse Alexa Flour Plus 800 (Invitrogen A32730). Protein bands were detected using the Odyssey CLx infrared imaging system (LICOR Biosciences).

### Immunofluorescence

Cells plated on coverslips were fixed in 10% neutral buffered formalin for 10 minutes at room temperature, washed three times with PBS, and permeabilized with .1% Triton X-100 for 10 min at room temperature. When MitoTracker staining was performed, cells plated on coverslips were stained with 100 nM MitoTracker (Thermo Fisher M22462) at 37°C for 15 minutes prior to fixation. Following permeabilization, coverslips were blocked for one hour at room temperature in blocking solution (Aqua block buffer, Abcam ab166952) and then transferred to a carrier solution (Aqua block) containing diluted primary antibodies: GOT2 (Sigma-Aldrich HPA018139), GOT2 (Thermo Fisher PA5-77990), COX IV (Cell Signaling Technology 11967S), COX2 (Abcam ab15191), His (R&D Systems MAB050-100). Coverslips were incubated with the primary antibody at 4°C overnight and then washed five times for 5 minutes each in PBS following which, secondary Alexa-flour conjugated antibodies diluted in the same carrier solution (1:400) were added to the coverslips for one hour at room temperature. After the secondary antibody incubation, coverslips were washed five times for five minutes each in PBS and mounted with Vectashield mounting media containing DAPI (Vector Laboratories H-1500). Images were captured on a Zeiss LSM 880 laser-scanning inverted confocal microscope in the OHSU Advanced Light Microscopy Shared Resource, and a 40×/ 1.1 NA water objective or 63x/1.4 NA oil objective was used to image the samples.

### Immunohistochemistry

Mice were anesthetized and euthanized according to institutional guidelines. Pancreatic tumors were excised carefully and fixed overnight in 10% phosphate-buffered formalin. Tissue samples were paraffin embedded and sectioned by the OHSU Histopathology Shared Resource. Human PDAC tissue sections from formalin-fixed, paraffin-embedded blocks were obtained from the OPTR. In brief, tissue sections were de-paraffinized and rehydrated through an ethanol series and ultimately in PBS. Following antigen retrieval, tissue samples were blocked for 1 hour at room temperature in blocking solution (8% BSA solution) and then transferred to a carrier solution (8% BSA solution) containing diluted antibodies: GOT2 (Sigma-Aldrich HPA018139), GOT2 (Thermo Fisher PA5-77990), COX IV (Cell Signaling Technology 11967S), COX2 (Abcam ab15191), CD3 (Abcam ab5690), CD4 D7D2Z (Cell Signaling Technology 25229S), CD8 (Abcam ab203035), Granzyme B (Abcam ab4059), F4/80 (Cell Signaling Technology 70076T), Arginase-1 (Sigma-Aldrich ABS535)). Sections were incubated overnight at 4°C and then washed five times for 5 minutes each in PBS. For fluorescence imaging, secondary Alexa-flour conjugated antibodies diluted in the same carrier solution (1:400) were added to the sections for one hour at room temperature. Sections were then washed five times for five minutes each in PBS and were mounted with Vectashield mounting media containing DAPI. For DAB chromogen imaging, sections were stained with primary antibody as described above, then the samples were incubated in polymeric horseradish peroxidase (HRP) conjugated secondary antibody (Leica PV6121) for one hour followed by 5 five-minute 1xTBST washes. HRP was detected using DAB chromogen (3,3′-Diaminobenzidine) solution (BioCare Medical BDB2004) prepared per manufacturer instructions. Tissues were exposed to chromogen solution until a brown precipitate was detected produced from oxidized DAB where secondary poly-HRP antibody is located. As soon as DAB chromogen is detected the tissue-slides were washed in diH2O, counterstained in hematoxylin, dehydrated and cleared for mounting. Stained tissue sections were scanned on a Leica Biosystems Ariol digital fluorescence scanner or Leica Biosystems Aperio brightfield digital scanner. Quantification was performed for single stains using QuPath quantitative pathology and bioimage analysis software v0.2.3. For co-stains (CD8/GRZB and F4/80/ARG1), manual counting was performed on at least 10 high-powered fields per tumor sample.

### Proliferation assays

PDAC cells were seeded into 96-well plates at 2 × 10^3^ cells per well in DMEM containing 10% FBS. Cells were treated as indicated in the manuscript text with 100 nM GW501516 (Cayman Chemical 10004272) at the time of cell seeding or 5mg/mL doxycycline (Sigma-Aldrich D9891) 48 hours prior to cell seeding. GW501516 and doxycycline treatments were both replenished every 48 hours for extended time points. After 72 hours or at the time points indicated in the manuscript, cells were lysed with CellTiter-Glo Luminescent Cell Viability Assay reagent (Promega) and luminescence was read using a GloMax plate reader.

### Chromatin immunoprecipitation

Chromatin immunoprecipitation was performed as described previously^5^. Briefly, PDAC cells were fixed in 1% formaldehyde, and nuclei were isolated and lysed in buffer containing 1% SDS, 10 mM EDTA, 50 mM Tris-HCl pH 8.0, and protease inhibitors, and sheared with a Diagenode Bioruptor to chromatin fragment sizes of 200–1000 base pairs. Chromatin was immunoprecipitated with antibodies to PPARδ (Abcam ab178866), or acetylated histone H3K9 (Cell Signaling Technology 9649). PPARδ binding or histone acetylation at known PPARδ target gene promoter regions was assessed by ChIP-qPCR and enrichment values were normalized to a control intergenic region of the genome. The following primer sequences were used: *Pparg* F: tgatgttgctgcaagggatg, R: agggttctatgctgaaggttct, *Tgfb1* F: ggtgctcgctttgtacaaca, R: gggtaatttcctcccggtga, *Cxcl10* F: gcatccctgagagaatcagc, R: gccaaatttagccagatcca, Intergenic A F: gacttcttcaccccacatgc, R: acagaggaaacagaaatggct, *Ptgs2* F: gggactcctcaggctcag, R: aagtgggttgcagttcctca, *Angptl4* F: tcagcctaccagggagagaa, R: ggaggaaagggcgtacaaat, *Cpt1a* F: ccaaacaaccaaacaaacca, R: cggcagacctaggacaacat, *Il6* F: gttctctgggaaatcgtgga, R: tcccaacctccactcaaaac, *Hmgcs2* F: ggccagagaaatgtttgagc, R: acctgcaacagcctctttgt, Intergenic B F: tggtgcttcttggtcaatca, R: aggacaaaacagcaaccaaca.

### Gene expression analysis by qPCR

The isolated total RNA (1 μg) was reverse-transcribed to produce cDNA using iScript Reverse Transcription Supermix kit (Bio-Rad). Real-time PCR was performed using SYBR Green supermix (Bio-Rad). The cDNA sequences for specific gene targets were obtained from the human genome assembly (http://genome.ucsc.edu) and gene specific primer pairs were designed using the Primer3 program (http://frodo.wi.mit.edu/primer3/primer3_code.html). Relative gene expression was normalized using the 36B4 housekeeping gene. The following primer sequences were used: human and mouse 36B4 (*RPLP0*): F:5’-GTGCTGATGGGCAAGAAC-3’; R:5’-AGGTCCTCCTTGGTGAAC-3’; human *PTGS2*: F: 5’-CTGGCGCTCAGCCATACAG-3’; R:5’CGCACTTATACTGGTCAAATCCC-3’; human *PDK4*: F:5’-AGAAAAGCCCAGATGACCAGA-3’; R:5’TGGTTCATCAGCATCCGAGT-3’; human *CPT1A*: F:5’-CTGTGGCCTTTCAGTTCACG-3’; R:5’-CCACCACGATAAGCCAACTG-3’; human *TGFB1*: F:5’-GGCCTTTCCTGCTTCTCAT-3’; R:5’-CAGAAGTTGGCATGGTAGCC-3’; human *CXCL1*: F:5’-AGGGAATTCACCCCAAGAAC-3’; R:5’-ACTATGGGGGATGCAGGATT-3’; human *CXCL9*: F:5’-ATTGGAGTGCAAGGAACCCC-3’; R:5’-ATTTTCTCGCAGGAAGGGCT-3’; human *CXCL10*: F:5’-GTGGCATTCAAGGAGTACCTC-3’; R:5’-TGATGGCCTTCGATTCTGGATT-3’; mouse *Ptgs2*: F:5’-TGAGTGGGGTGATGAGCAAC-3’; R:5’-TTCAGAGGCAATGCGGTTCT-3’; mouse *Pdk4*: F:5’-TGAACACTCCTTCGGTGCAG-3’; R:5’-GTCCACTGTGCAGGTGTCTT-3’; mouse *Ppargc1a*: F:5’-GCTTGACTGGCGTCATTCGG-3’; R:5’-TGTTCGCAGGCTCATTGTTG-3’; mouse *Tgfb1*: F:5’-CACTCCCGTGGCTTCTAGTG-3’; R:5’-GTTGTACAAAGCGAGCACCG-3’; mouse *Cxcl1*: F:5’-TGGCTGGGATTCACCTCAAG-3’; R:5’-CCGTTACTTGGGGACACCTT-3’; mouse *Cxcl9*: F:5’-AACGTTGTCCACCTCCCTTC-3’; R:5’-CACAGGCTTTGGCTAGTCGT-3’; mouse *Cxcl10*: F:5’-CAAGCCATGGTCCTGAGACA-3’; and R:5’-TGAGCTAGGGAGGACAAGGA-3’.

### Metascape analysis

PDA TCGA Firehose Legacy data base provides mRNA expression data for co-expression analysis accessible through cBioportal. The data set includes Spearman’s correlation analysis and P-values for each gene comparison. The data set was used to identify genes negatively correlated with GOT2 expression in PDA patients. A list of genes with Spearman’s correlation value of equal or less than −0.25 and a P-value of less than 0.01 was generated. The list of genes was submitted to online bioinformatics tool Metascape for identification of enriched gene ontology clusters in the data set. The output from Metascape analysis was graphed using GraphPad Prism.

### Orthotopic PDAC modeling

The orthotopic transplant method used here was described previously^6^. In brief, 8- to 10-week-old wild-type male C57BL/6J (for FC1245) or B6129SF1/J (for 688M) mice were orthotopically transplanted as described previously with 5 × 10^3^ FC1245 cells or 8 × 10^4^ 688M cells in 50% Matrigel (Corning 356231), 50% DMEM. For experiments with 688M cells harboring VP16-PPARD, 6 × 10^4^ 688M cells were used. For pharmacologic activation of PPARδ, mice were treated with vehicle (5% PEG-400, 5% Tween-80 in diH2O) or with 4 mg/kg GW501516 in vehicle by intraperitoneal injection once daily. Vehicle was created and autoclaved before use. GW501516 was created in 10 mM stock in DMSO and stored in 250 μl aliquots at −20°C (one for each day of treatment). On the day of treatment, a vial was thawed, diluted 1:10 in vehicle, and mice were dosed at 4 mg/kg. For T cell neutralization experiments, mice received intraperitoneal injection of 0.2 mg of αCD8 (2.43), αCD4 (GK1.5), or an IgG2b isotype control (LTF-2) diluted in 100 μl sterile PBS. Antibodies were purchased from BioXcell and were administered beginning 2 days pre-implantation with 6 × 10^4^ 688M cells and every 4 days thereafter until euthanasia, as previously described^7^. Mice were euthanized when control animals were moribund, and tumors were excised, weighed, and immediately fixed in formalin.

### Long chain fatty acid binding site prediction

The arachidonic acid binding site on the human GOT2 surface is predicted using the molecular modeling technique. Druggable hotspots identification has long been used to predict and explore the allosteric pockets that accommodate substrate and drug-like molecules^8,9^. A similar approach is taken to identify a plausible arachidonic acid binding site by probing the GOT2 3-D protein structure^10^ (PDBID:5AX8). The protein structure was prepared using the protein prep tool of Maestro-2014-3 (Schrödinger, LLC, New York, NY, 2013). Arachidonic acid is a 20 carbon long-chain fatty acid (LCFA) with greasy carbons and a carboxylate group. The available structural information suggests that the binding pocket must be hydrophobic with the positively charged residues to accommodate LCFA^11,12^.

The SiteMap^13^ calculation accounts for the prediction of pockets, characterized by cavity volume, chemical, and physical properties as that of known druggable sites. Five sites were predicted on the GOT2 structure, and these sites had a site score of > 0.8, composed of hydrophobic, hydrogen bond acceptor, and donor volumes. The top-ranked site-1 is a catalytic site, and site-2 to 5 are allosteric. Arachidonic acid docked against all the predicted sites. The Induced-Fit docking protocol^14^ adopted here allows both the ligand and the surrounding residues of protein to be flexible. A total of five docking runs were performed on the predicted site. The docking grid boxes are defined based on the residues suggested by the SiteMap analysis (Site-1: N215, H210; Site-2: N270, F239; Site-3: A260, W226, H373, G385, Q390; Site-4: R337, G254; Site-5: N332, D93). The site-2 ~25 Å away from the catalytic site resulted in a binding pose with favorable energy and interaction complementarity between the protein and ligand. Compared to other sites, Site-2 has increased hydrophobic volume, which may recognize LCFA like arachidonic acid. Triple mutants K234A/K296/R303 were proposed to validate the predicted binding pose. K234 interacts with the carboxylate group of LCFA. K296, which is in proximity to making ionic interaction (in dynamics) and perturbation of the positive charge to neutral alanine residues, prevents the charged interaction. From the docking pose, R303 is making the hydrophobic interaction with the lipid tail of arachidonic acid. R303A mutation reduces the hydrophobic interaction by the sidechain of arginine. The proposed triple mutations have the potential to abolish the arachidonic acid binding.

### Fatty acid binding assay

Reactions were carried out in binding buffer (0.003% digitonin in 1X PBS) containing 1 μM of purified human GOT2 protein (AA30-430) and 0.5 μci/ml [3H]-arachidonic Acid. After incubation for 1hr at 4°C, the mixture was incubated with pre-equilibrated of TALON Metal Affinity Resin (Takara, 635502) at 4°C for 1 h, then loaded onto a column and washed with binding buffer, then binding buffer with 0.01% BSA, and binding buffer again. The protein-bound [3H]-arachidonic was eluted with elution buffer (50 mM sodium phosphate, 300 mM sodium chloride, 150 mM imidazole; pH 7.4.) and quantified by scintillation counting. For competition experiments with unlabeled lipids, the assays were carried out in the presence of ethanol containing the indicated unlabeled sterol (0 –1 mM).

### Luciferase assay

PPRE x3-TK-Luc (PPAR response element driving luciferase) plasmid #1015 was purchased from Addgene and the Renilla plasmid (pRL-SV40) was generously provided by Dr. Ellen Langer (OHSU). Cells were transfected with 2.5 μg PPRE x3-TK-Luc, 15 ng pRL-SV40, and 4 μl Lipofectamine 2000 in 6-well plates. Briefly, cells were plated at 1 × 10^6^ per well of a 6-well plate and allowed to adhere overnight. Plasmids were combined in 150 μl Opti-MEM while lipofectamine 2000 was combined in a separate tube with 150 μl Opti-MEM. After 5 mins the tubes were combined. 300 μl of the mixture was added, in a dropwise manner, to 700 μl of Opti-MEM on each well for transfection. The cells were incubated overnight at 37°C, collected, counted, and replated to white-walled 96-well plates in triplicates. 24 hours later a dual luciferase assay was completed, following the manufacturer’s instructions: Dual-Luciferase Reporter Assay System (Promega E1910). Briefly, cells were lysed in white-walled 96-well plates with 20 μl 1X Passive Lysis Buffer and shaken on a room temp shaker. 100 μl LARII was added to each well and luminescence was measured over 5 seconds. 100 μl of Stop and Glo were then added and Renilla activity was measured with luminescence over 5 seconds. Activity was calculated by normalizing luciferase signal to renilla for each well.

### PPARδ transcription factor activity assay

Nuclear lysates were prepared using a detergent-free fractionation protocol. Cells were scraped and collected from 10 cm dishes, washed with PBS, pelleted (450 x g 5mins), resuspended in PBS and 1/5 of the volume was reserved for whole cell lysis in RIPA (Amresco N653-100mL) + cOmplete EDTA-free Protease inhibitor cocktail (Sigma-Aldrich 11836170001). The remaining 4/5 of cell suspension was centrifuged (450 x g 5mins), PBS removed and cells were lysed on ice for 15 mins in Lysis buffer (5x of cell pellet volume). Lysis Buffer: 10mM HEPES pH 7.9, 1.5mM MgCl2, 10mM KCl with 1mM DTT and EDTA-Free cOmplete mini protease inhibitor cocktail. Lysates were centrifuged (450 x g 5mins), supernatant decanted, lysis buffer added (2x cell volume) and suspensions ground on ice with a plastic homogenizer 10x in 1.5mL Eppendorf tubes. Lysates were centrifuged (10,000 x g 20mins) and supernatant collected as cytosolic fraction. Remaining pellet was washed with 200 μl lysis buffer (10,000 x g 5mins), supernatant decanted and extraction buffer added (2/3x cell pellet volume). Extraction buffer: 20mM HEPES pH 7.9, 1.5mM MgCl2, 0.42M NaCl, 0.2uM EDTA, 25% glycerol (V/V), 1mM DTT and cOmplete mini EDTA-free protease inhibitor cocktail. Nuclei were ground with plastic homogenizer in 1.5mL Eppendorf tubes 20x, and incubated at 4°C with gentle shaking for 10 mins. Samples were centrifuged (20,000 x g 5mins) and supernatant transferred to cold Eppendorf tubes, as nuclear fraction. Lysates were measured with BCA and an equal protein amount was added per sample for each well. Manufacturer’s instructions were followed for the PPAR delta transcription factor kit (Abcam ab133106). Briefly CTFB was prepared and added to blank and NSB wells, nuclear lysates were added to each sample well containing immobilized PPRE-containing DNA and the plate was incubated overnight at 4°C without agitation. The next day the wells were washed 5x in 1X wash buffer and incubated in PPARdelta primary antibody (1:100) for 1hr at room temperature in the dark, without agitation. Wells were washed 5x in 1X wash buffer and incubated in goat anti-rabbit HRP conjugate (1:100) for 1hr at room temperature in the dark without agitation. Wells were washed 5x in 1X wash buffer and 100 μl developing solution was added to each well. Plate was incubated for 15-45 minutes on a room temperature shaker, in the dark, until color developed. 100 μl stop solution was added to wells and the absorbance at 450nm was taken.

### Nuclear fatty acid uptake assay

MiaPaca2 ctrl and sh27 cells were plated at 5 × 10^5^ in a 6 well plate and allowed to adhere overnight. Media was changed to 0% FBS DMEM and the cells were incubated for 24hrs. The media was changed to 0.5% Fatty-Acid Free BSA DMEM with either chloroform (ctrl) or 2.5 μM NBD-arachidonic acid (Avanti Polar Lipids 810106C). Media was made before being added to cells, heated to 37°C and vortexed until fatty acid was completely in solution. Cells were incubated at 37°C for durations indicated in the manuscript and collected and fractionated using the Detergent Free Method described above (PPARδ transcription factor activity assay). Nuclear lysates were placed in a white-walled 96-well plate and fluorescence was measured at 480 nm excitation and 540 nm emission. Lysate concentration was measured using a BCA kit. FC1245 cells were plated 5 × 10^5^ per well and treated as described above, but treatment was reduced to 2 μM NBD-arachidonic acid for 15 minutes due to lipid toxicity in this cell line.

### Aspartate aminotransferase assay

AST Activity Assay Kit (Sigma-Aldrich MAK055) was used to determine aspartate aminotransferase activity per manufacturer instructions. Briefly, this assay determines the transfer of an amino group from aspartate to alpha-ketoglutarate in the generation of glutamate which produces a colorimetric product (450 nm) that is proportional to aspartate aminotransferase activity in the sample. For this assay Pa-Tu-8988T cells with stable expression of doxycycline inducible GOT2 shRNA were transiently transfected with wtGOT2 and tmGOT2. After 48hrs these cells were exposed to doxycycline for 48hrs to knockdown endogenous GOT2 in cells with GOT2 shRNA. Cells were seeded at 5 × 10^6^ and collected via trypsin disassociation after cells were adhered. The cells were then resuspended in 1ml of ice-cold 1X PBS and 200 μl (1 × 10^6^ cells) were collected for AST assay and 800 μl (4 × 10^6^ cells) were collected for protein concentration estimation and Western blot protein expression analysis. Using AST assay kit buffers, cells were lysed to obtain a supernatant which was combined with the kit reagent master mix to detect glutamate in a colorimetric reaction. The samples were read every 5 minutes for 30 minutes. AST activity and concentration in the samples were determined using instructions from the manufacturer.

### Free fatty acid measurements

Samples were subjected to an LCMS analysis to detect and quantify levels of free fatty acids in sample extracts. A fatty acid extraction was carried out on each sample using 100% methanol as the homogenization solvent. Whole cell pellets (1 × 10^6^ cells/sample) were lysed with 1000 μL of methanol and ~100 μL of zircon beads (0.5 mm). Manual disruption with a p1000 pipette tip was performed to assist initial pellet suspension in extraction buffer. The methanol extracts were centrifuged (21,000g x 3 min) and transferred to glass LCMS inserts for analysis. The LC column was a WatersTM BEH-C18 (2.1 ×100 mm, 1.7 μm) coupled to a Dionex Ultimate 3000TM system and the column oven temperature was set to 25°C for the gradient elution. The flow rate was 0.1 mL/min and used the following buffers; A) water with 0.1% formic acid and B) acetonitrile with 0.1% formic acid. The gradient profile was as follows; 60-99%B from 0-6 min, hold at 99%B from 6-10 min, 99-60%B from 10-11 min, hold at 60%B from 11-15 min. Injection volume was set to 1 μl for all analyses (15 min total run time per injection).

MS analyses were carried out by coupling the LC system to a Thermo Q Exactive HFTM mass spectrometer operating in heated electrospray ionization mode (HESI). Data acquisition was 10 min with a negative mode full MS scan (profile mode) and one microscan, with an AGC target of 3e6 and a maximum IT of 100 ms at 120,000 resolution, with a scan range from 160-400 m/z. Spray voltage was 3.5kV and capillary temperature was set to 320°C with a sheath gas rate of 35, aux gas of 10, and max spray current of 100 μA. The acquisition order of samples and standard curve points was randomized, with blank matrix controls before and after each standard curve point to assess carry over (none detected). The resulting free fatty acid peaks were quantified by measuring the relative intensities (peak heights) of the high resolution extracted ion chromatogram (XIC) for each fatty acid across the samples and external standard curve samples ranging from 10 μg/mL to 100 ng/mL. All fatty acids were detected as the negative mode [M-H] ion and retention times of the fatty acids were defined using a cocktail of authentic standards. For each XIC, the theoretical m/z of each fatty acid (± 5 ppm) was used to extract the peak height (24 sec retention time window, 12 sec retention time tolerance) as follows: Lauric acid (199.1704 m/z, 2.3 min), Myristic acid (227.2017 m/z, 3.1 min), Palmitoleic acid (253.2173 m/z, 3.4 min), Palmitic acid (255.2330 m/z, 4.1 min), Oleic acid (281.2486 m/z, 4.4 min), Stearic acid (283.2643 m/z, 5.1 min), Arachidic acid (311.2956 m/z, 6.0 min), Nervonic acid (365.3425 m/z, 6.9 min), Lignoceric acid (367.3582 m/z, 7.5 min). The resulting standard curve points (in duplicate) were fit to a linear regression (GraphPad Prism8), and this equation was used to interpolate the concentration of fatty acids in the sample extracts, as prepared.

### Statistical Analysis

All statistical analyses were performed using GraphPad Prism 9.0 Software (Graph Pad Software Inc.).

## Acknowledgements

We thank all members of the Sherman lab as well as Drs. Sara Courtneidge and Amy Moran for helpful discussion of this work. This study was supported by a postdoctoral fellowship from the OHSU Fellowship for Diversity in Research (to J.A.), a graduate student fellowship from the Knight Cancer Institute Cancer Center Support Grant P30 CA069533 (to H.S.-C.), NIH grants R00 CA188259 and R01 CA229580 (to M.H.S.) including CA229580-S1 Research Supplement to Promote Diversity in Health-Related Research (awarded to M.H.S. in support of J.A.), American Cancer Society grant RSG-18-142-01-CSM (to M.H.S.), and an OHSU Faculty Innovation Fund Award (to M.H.S.). We thank members of the OHSU Histopathology Shared Resource, Medicinal Chemistry Core, and Advanced Light Microscopy Shared Resource for supporting this study, as well as Dr. Drew Jones and Leonard Ash from the NYU Langone Health Metabolomics Laboratory.

## Author contributions

H.S.-C., J.A. and M.H.S. conceived the study and designed the experiments; H.S.-C. and J.A. generated all cell lines and performed the majority of the experiments including all mouse studies; C.O. generated the VP16-PPARD and GOT2 constructs, aided in cell line generation, and performed *in vitro* studies; X.X. performed the competitive fatty acid binding assays with guidance from P.T.; S.N. performed the GOT2 docking studies and computational modeling of arachidonic acid binding; S.B. performed immunohistochemistry on PDAC patient samples; all authors contributed to data analysis; H.S.-C., J.A. and M.H.S. wrote the paper with input from all authors.

## Additional information

Supplementary Information is available for this paper. Correspondence and requests for materials should be addressed to M.H.S.

---

**Extended Data Figure 1:**
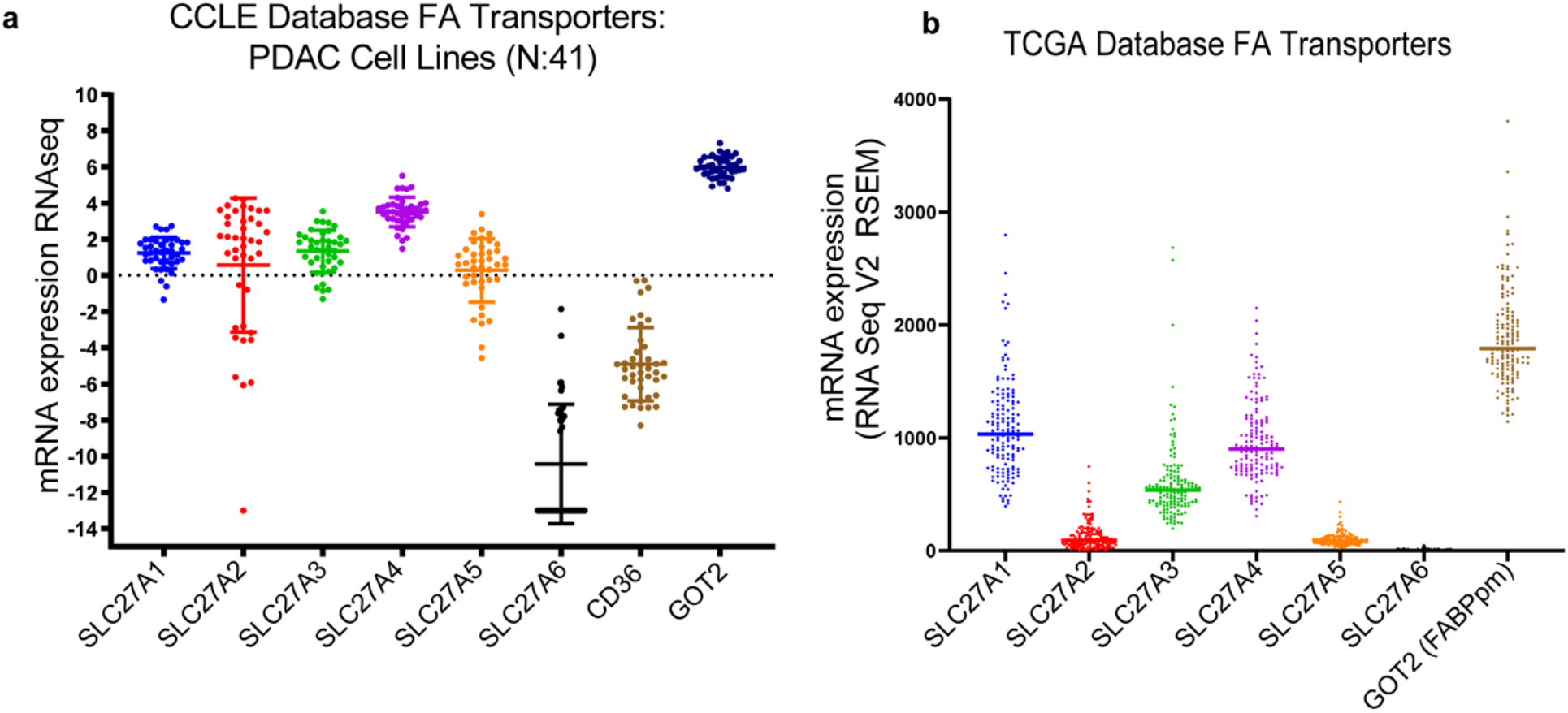
GOT2 is upregulated in human PDAC. **a,** Relative expression of the indicated fatty acid transporters and trafficking factors among 41 human PDAC cell lines in the Broad Institute Cancer Cell Line Encyclopedia database. **b,** Relative expression of the indicated fatty acid transporters and trafficking factors in human PDAC in The Cancer Genome Atlas database.

**Extended Data Figure 2:**
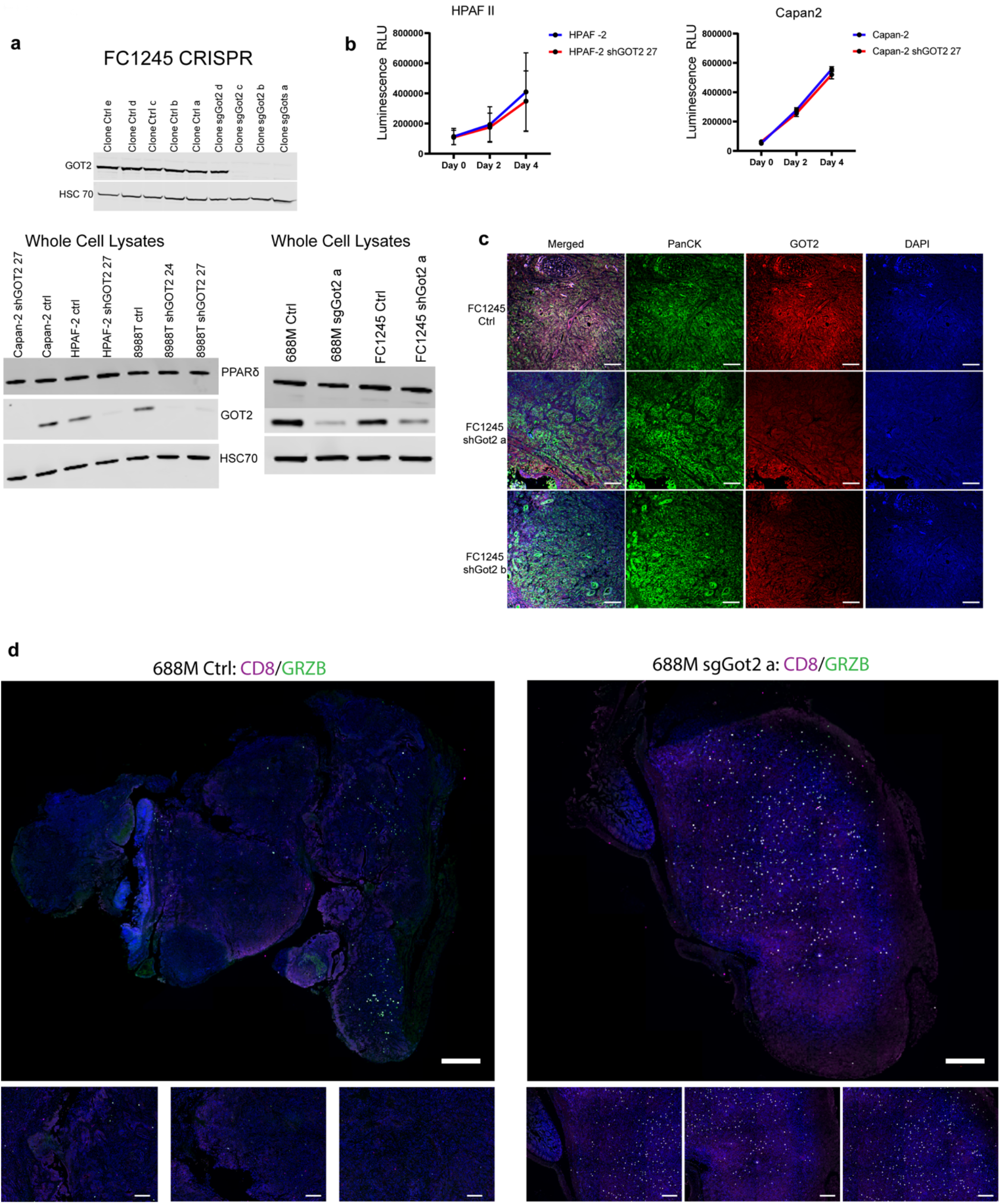
PDAC cells maintain proliferative capacity with loss of GOT2 expression. **a,** Western blots indicating GOT2 levels in control and GOT2 loss-of-function PDAC lines. **b,** Viable cell measurements in the indicated PDAC lines. Data are presented as mean ± s.e.m. from biological triplicates. **c,** Immunohistochemical staining of FC1245 tumors at experimental endpoint, representative of n = 3 per cohort. Scale bar = 50 μm. **d,** Immunohistochemical staining of 688M tumors at experimental endpoint, representative of n = 3-5 per cohort, at low magnification to display tissue-wide staining patterns. Upper images: scale bar = 1 mm, lower images: scale bar = 500 μm.

**Extended Data Figure 3:**
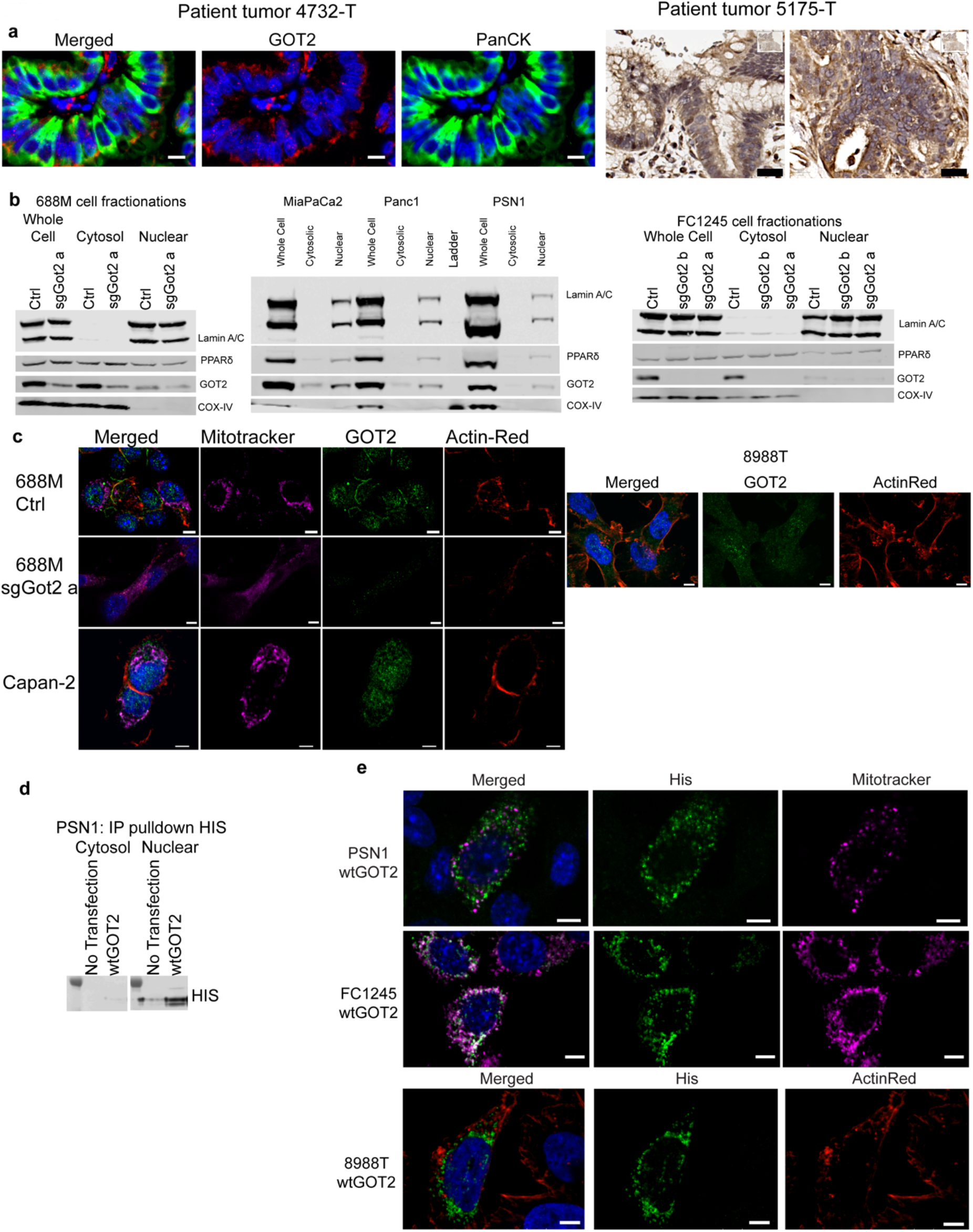
A pool of GOT2 protein localizes to the nucleus in PDAC cells. **a,** Immunohistochemical staining for GOT2 or GOT2 and panCK in human PDAC (representative of n = 5), showing additional samples to complement Figure 3b. Fluorescent images: scale bar = 2 μm, brightfield image: scale bar = 20 μm. **b,** Western blots in PDAC cell lines indicating GOT2 protein levels in the indicated cellular fractions. Lamin A/C is a loading control for nuclei, COX-IV is a loading control for cytoplasm and indicates an absence of mitochondrial protein in the nuclear fraction. **c,** Immunofluorescent staining of the indicated PDAC cell lines for endogenous GOT2. Mitotracker indicates mitochondria, Actin-Red indicates F-actin, and nuclei are counterstained with DAPI. Scale bar = 5 μm. **d,** Immunoprecipitation of transiently transfected, His-tagged GOT2 from the indicated cellular fractions in PSN-1 human PDAC cells. **e,** Immunofluorescent staining of the indicated PDAC cell lines for transiently transfected, His-tagged GOT2. Scale bar = 5 μm.

**Extended Data Figure 4:**
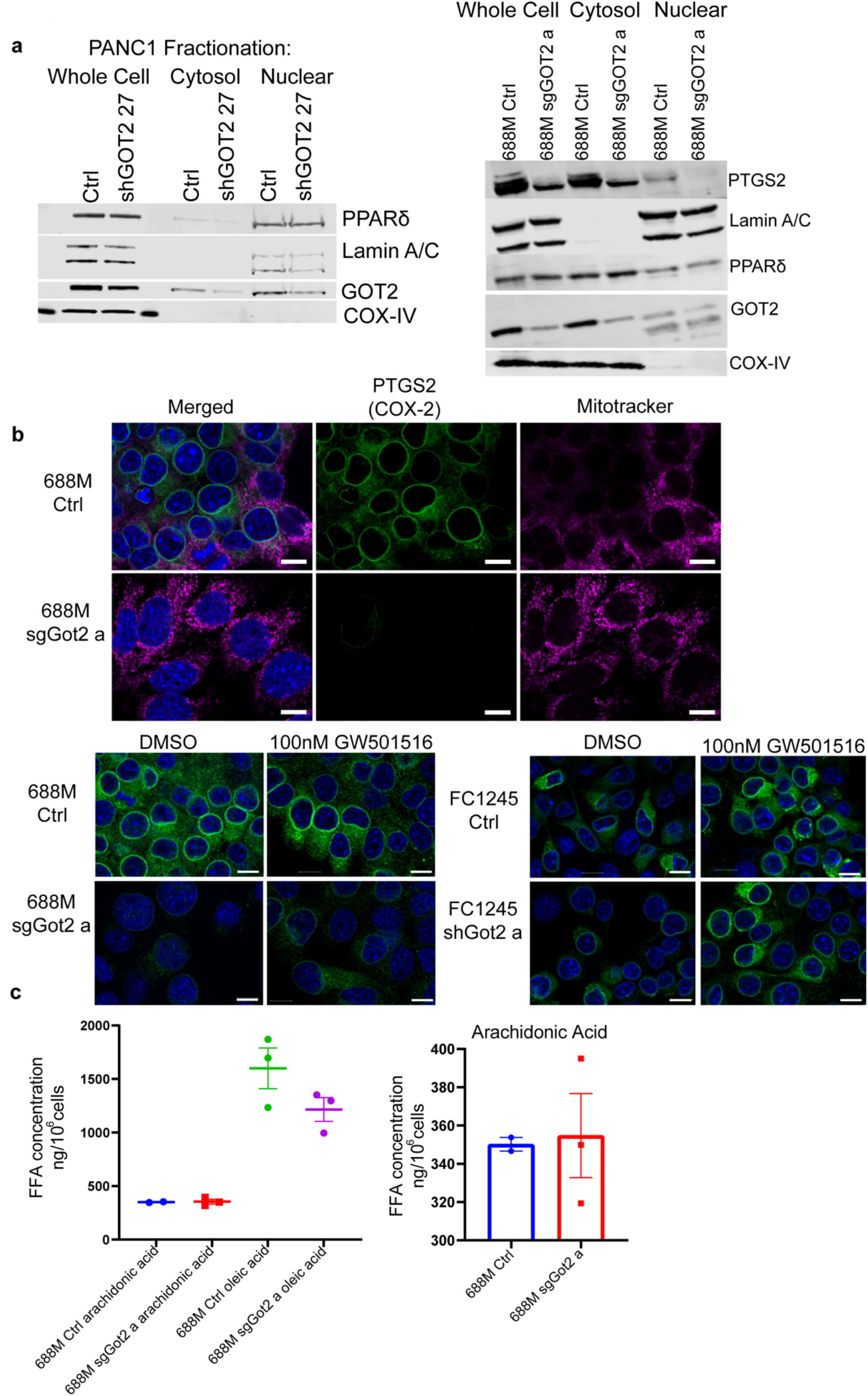
GOT2 positively regulates PPARδ activity and target gene expression. **a,** Western blots indicating levels of GOT2, PPARδ, and PPARδ target PTGS2/COX2 in the indicated PDAC lines. **b,** Immunofluorescent staining of ctrl and sgGot2 688M cells, as well as ctrl and shGot2 FC1245 cells, for PTGS2/COX2 with or without 100 nM GW501516 treatment. Scale bar = 10 μm. **c,** Quantification of fatty acid levels in ctrl and sgGot2 688M cells, measured in whole cells by LCMS.

**Extended Data Figure 5:**
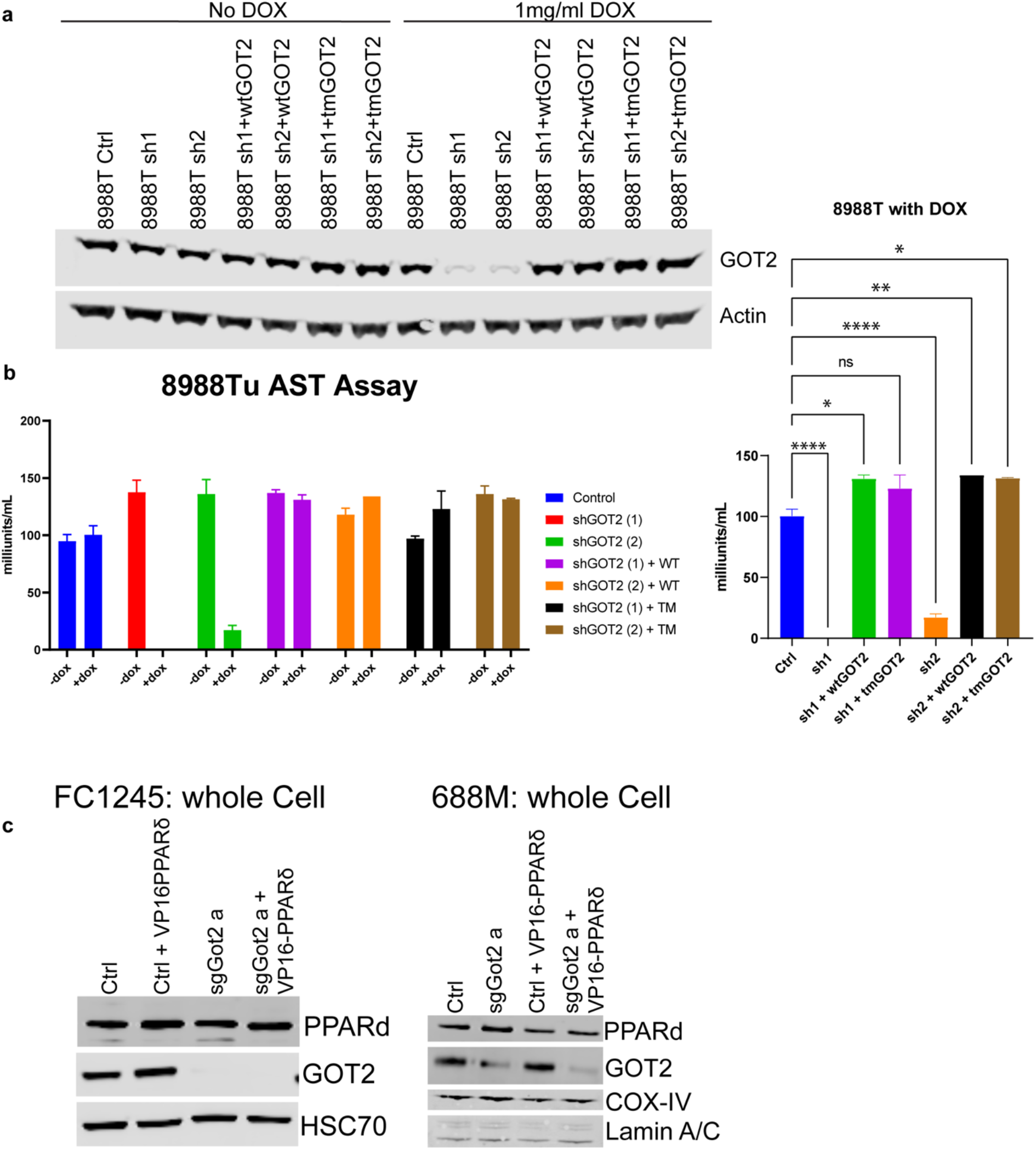
GOT2 fatty acid binding and PPARδ activation support tumor growth. **a,** Western blots indicating GOT2 levels in doxycycline-inducible GOT2 knockdown 8988T cells, reconstituted with wtGOT2 or tmGOT2. **b,** Aspartate aminotransferase activity assay (also known as glutamate-oxaloacetate transaminase activity assay) on the cells indicated in **a**. Data are plotted as mean ± s.e.m. from biological triplicates. *p < 0.05, **p < 0.01, ****p < 0.0001 by one-way ANOVA. **c,** Western blots indicating GOT2 and PPARδ expression in the indicated stable cell lines.

